# Characterisation of *Staphylococcus aureus* lipids by nanoelectrospray ionisation tandem mass spectrometry (nESI-MS/MS)

**DOI:** 10.1101/593483

**Authors:** Simon A. Young, Andrew P. Desbois, Peter J. Coote, Terry K. Smith

**Author notes:** Corresponding author: (SY).

## Abstract

*Staphylococcus aureus* is a major opportunistic pathogen that is exposed to antimicrobial innate immune effectors and antibiotics that can disrupt its cell membrane. An understanding of *S. aureus* lipid composition and its role in defending the cell against membrane-disrupting agents is of fundamental importance. Common methods for characterising lipid profiles suffer shortcomings such as low sensitivity of detection and inferior resolution of the positional assignments of fatty acid chains in lipids. This present study developed a rapid and sensitive nano-electrospray ionisation tandem mass spectrometry (nESI-MS/MS) method to characterise the lipid composition of three commonly studied *S. aureus* isolates: Newman, Mu50 and BB270. Confirming previous studies, nESI-MS/MS revealed that phosphatidylglycerols were most abundant in *S. aureus* membranes, while diglucosyldiacylglycerols and lysyl-phosphatidylglycerols were also detected. Positional assignments for individual fatty acid chains within these lipids were also determined. Concomitantly, gas chromatography mass spectrometry of the fatty acids validated the molecular characterization and showed the principal species present in each strain were predominately anteiso- and iso-branched chain fatty acids. Though the fatty acid and lipid profiles were similar between the *S. aureus* strains, this method was sufficiently sensitive to distinguish minor differences in lipid composition. In conclusion, this nESI-MS/MS methodology can characterise the role of lipids in antimicrobial resistance, and may even be applied to the rapid diagnosis of drug-resistant strains in the clinic.

## Introduction

As a Gram-positive bacterium found in the respiratory tract and on the skin of healthy humans, *Staphylococcus aureus* is a major opportunistic pathogen responsible for a range of community-acquired and nosocomial topical or systemic infections [1]. These infections are treated with various classes of antibiotics, including agents that exert their antimicrobial action by disrupting the bacterial cell membrane, such as daptomycin [2]. Moreover, many innate immunity defence effectors that act during the early stages of infection, such as free fatty acids and cationic antimicrobial peptides (AMPs), also exert antimicrobial action through targeting the integrity and functioning of the cell membrane [3,4,5].

The major components of cell membranes are phospholipids, which form a semipermeable barrier to maintain cell homeostasis and prevent the entry of harmful substances. Phospholipid-containing membranes can permit bacteria to persist in the face of external hazards such as osmotic stress or extremes of pH, and they play an integral role in infection, acting as a barrier to antibiotics and host defense mechanisms [6]. Far from being static structures, bacteria constantly modify the lipid composition of their membranes in response to changes in the physiological environment, such as temperature, osmolarity, salinity and pH [7]. Thus, it is of fundamental and clinical importance to gain a detailed understanding of the composition of the *S. aureus* membrane and the role that changes in lipid composition might render the cells less or more susceptible to the action of membrane-disrupting antimicrobial agents.

The membranes of *S. aureus* have a preponderance of three phospholipids, phosphatidylglycerol (PG), lysyl-phosphatidylglycerol (L-PG) and cardiolipin (CL), in addition to other non-polar, glycosylated or conjugated lipid species [8]. Various approaches have been used to characterise this lipid composition; most studies have utilised column chromatography and two-dimensional thin-layer chromatography (2D-TLC) to provide basic identification and chemical composition of lipid species [9,10]. Initially such characterisation was performed on protoplasts derived from late-logarithmic or stationary phase *S. aureus* cultures, thereby providing only a restricted view of the lipid composition. Long-term radiolabelling of logarithmically growing bacteria with either [^14^C]acetate or [2-^3^H]glycerol helped to provide a more complete picture of the lipid content when analysed by 2D-TLC [11]. This provided an estimate for the relative abundances of the different lipid species in the cell membranes, notably ~50% PG, ~20% diacylglycerol, ~10% L-PG, ~7% diglucosyldiacylglycerol, ~1% CL, ~1% monoglucosyldiacylglycerol and ~5% of the cell wall polymer lipoteichoic acid (LTA) [11]. Interestingly, CL content has been found to increase in stationary phase [12] and under conditions of high salt [13]. When transitioning from exponential growth to stationary phase CL content increases six-fold and this is accompanied by a corresponding reduction in PG content, though the relative abundance of L-PG remains constant [12]. Nevertheless, while 2D-TLC can provide accurate quantification of each lipid class, it requires large amounts of lipid extracts and does not provide specificity with regard to individual molecular species [14]. The application of targeted mass spectrometry methods has provided some insight into the chemical composition of specific *S. aureus* lipids [15,16], while gas chromatography mass spectrometry (GC-MS) has proved successful in determining the specific fatty acid composition of *S. aureus* following diverse genetic, chemical and environmental manipulations [17,18,19].

Nevertheless, a need exists for a rapid, sensitive and accurate quantitative method to characterise in detail the lipid composition of the *S. aureus* cell membrane. The recent applications of two distinct liquid chromatographic (LC) MS approaches, reversed-phase LC-MS [20] and hydrophilic interaction LC-ion mobility-MS [21], have addressed some of the limitations in earlier methodologies. Both of these studies describe similar, but not identical *S. aureus* lipidomes, but these remain incomplete, as the authors did not obtain positional assignments for the individual fatty acid chains to enable a full characterisation of each lipid species. While it is common to use high-performance liquid chromatography (HPLC) methodologies to separate lipids prior to MS detection, this is not essential because of the relatively simple nature and lack of diversity of the *S. aureus* lipidome. In addition, LC-MS approaches are time consuming and require significant downstream data processing and analysis. To this end, we have utilised nanoelectrospray mass spectrometry (nESI-MS and nESI-MS/MS) in combination with a direct infusion non-targeted (shotgun) methodology to perform detailed characterisation of the lipid profile and fatty acid composition of the abundant lipids in *S. aureus* cell membranes. This approach (in addition to subsequent GC-MS analysis of fatty acid chains) provides a rapid and convenient way to sensitively compare membrane lipid compositions between *S. aureus* strains.

## Materials and methods

### *S. aureus* culture and lipid extraction

All reagents and culture media were purchased from Sigma-Aldrich Ltd. *S. aureus* Newman (methicillin-susceptible; MSSA), BB270 (methicillin-resistant; MRSA) and Mu50 (MRSA and vancomycin intermediate-resistant) were sourced and cultured in Müller-Hinton broth as described previously [22]. Briefly, each bacterial strain was prepared by inoculation of a colony from an agar plate into 10 ml Müller-Hinton broth and cultured at 37°C with shaking until reaching an OD_600nm_ = 1 (~1 × 10^9^ colony-forming units [cfu]). Bacterial cells were harvested (1000 × g, 10 min), washed in PBS before a repeat centrifugation and removal of the supernatant. Lipids were extracted according to the Bligh-Dyer method [23], dried under nitrogen, and stored at 4°C until analysis by mass spectrometry.

### Fatty acid methyl ester preparation and GC-MS analysis

The global fatty acid composition of the *S. aureus* strains was characterised and quantified by their conversion to fatty acid methyl esters (FAME) followed by GC-MS analysis. Briefly, duplicate aliquots of the lipid extracts equivalent to 1 × 10^8^ cfu were transferred to 2-ml glass vessels and dried under nitrogen gas. Fatty acids were released from intact lipids by base hydrolysis using 500 μl of concentrated ammonia and 500 μl of 50% propan-1-ol (in water), followed by incubation for 5 h at 50°C. After cooling to room temperature the samples were evaporated to dryness with nitrogen gas and then dried twice more after washing in 200 μl of methanol:water (1:1) to remove all traces of ammonia. The protonated fatty acids were then extracted by partitioning between 500 μl of 20 mM HCl and 500 μl of ether, before the aqueous phase was re-extracted with fresh ether (500 μl) and the combined ether phases were dried under nitrogen gas in a glass tube. The fatty acids were converted to methyl esters by adding diazomethane (3 × 20 μl aliquots) to the dried residue, while on ice. After 30 min the samples were allowed to warm to room temperature and left to evaporate to dryness in a fume hood. The dried FAME samples were dissolved in 20 μL dichloromethane and analysed by injection of 1 μL into a GC-MS (GC-6890N, MS detector-5973; Agilent Technologies) using a ZB-5 column (30 m × 25 mm × 25 mm; Phenomenex) operating a temperature program of 50°C for 10 min, followed by a rising gradient to 220°C at 5°C min^−1^, and held at 220°C for a further 15 min. Mass spectra were continuously acquired in the range of 50-500 atomic mass units (amu) and peak identification was performed by comparison of retention times and fragmentation patterns with a standard bacterial FAME mixture containing both odd and even fatty acid chains (Supelco 47080-U; Sigma-Aldrich Ltd).

### nESI-MS/MS analysis

Lipid extracts were analysed by nESI-MS and nESI-MS/MS following the method of Lilley *et al*. [24]. Briefly, total lipid extracts were dissolved in 100 μl of chloroform:methanol (1:2) and analysed with a triple quadrupole time of flight (QTOF) mass spectrometer equipped with a nanoelectrospray source (4000 QTrap; AB SCIEX). Each sample (15 μl) was delivered using a Nanomate interface (Advion, Inc.) in direct infusion mode (~125 nl min^−1^). The lipid extracts were analysed in both positive and negative ion modes using a capillary voltage of 1.25 kV. MS/MS scanning (precursor, daughter and neutral loss) were performed using nitrogen as the collision gas and each spectrum encompassed at least 50 repetitive scans. Tandem mass spectra were obtained with collision energies as follows: 20-35 V, neutral loss scanning of m/z 300 in positive ion mode to detect L-PG; 50 V, selected precursor ion (m/z 211, 225, 239, 253, 267, 281, 295, 309, 323) scanning in negative mode. MS/MS fragmentation/daughter ion scanning was performed in positive and negative modes with collision energies between 35–100 V. For each sample, data were acquired in two distinct scan windows (600–1000 m/z and 1000–1500 m/z) in both positive and negative modes. The assignment of individual lipid species was based upon a combination of survey, daughter and precursor scans, as well as submitting parent ion masses to the LIPID MAPS database at the Nature Lipidomics Gateway (http://www.lipidmaps.org).

## Results and discussion

### Fatty acid composition

GC-MS analysis of the FAME prepared from total lipid extracts from the three laboratory-cultured *S. aureus* strains, BB270, Newman and Mu50, revealed that the principal fatty acid species ranged from C14 to C20 in length (Fig 1A). In addition, the analysis confirmed the dominant presence of iso- and anteiso-branched chain fatty acids, common to Gram-positive bacteria [25]. All three strains showed similar profiles, although when cultured in Müller-Hinton broth the BB270 lab strain consistently had a reduced percentage (4.53 ± 0.25 %) of longer straight- and branched-chain fatty acids (specifically i-19:0, a-19:0, 19:0, 20:0) relative to the Newman (7.72 ± 0.47 %) and Mu50 (10.56 ± 0.27 %) strains. Grouping the fatty acids by structure as straight, anteiso- and iso-branched, confirmed that the strains have relatively similar proportions of each type of fatty acid (Fig 1B). However, calculating the specific ratios of branched-to straight-chain fatty acids revealed values of 4.57 ± 0.20 (BB270), 10.9 ± 0.36 (Newman) and 9.06 ± 0.23 (Mu50), indicating the BB270 strain to have a greater proportion of saturated straight-chain fatty acids. Notably this was due to a reduced proportion of anteiso-branched chain fatty acids, as the proportion of iso-branched chains remained similar between the strains (Fig 1B). Overall, the fatty acid composition of these strains closely matched those of other *S. aureus* strains cultured in Müller-Hinton broth [26].

**Fig 1.**
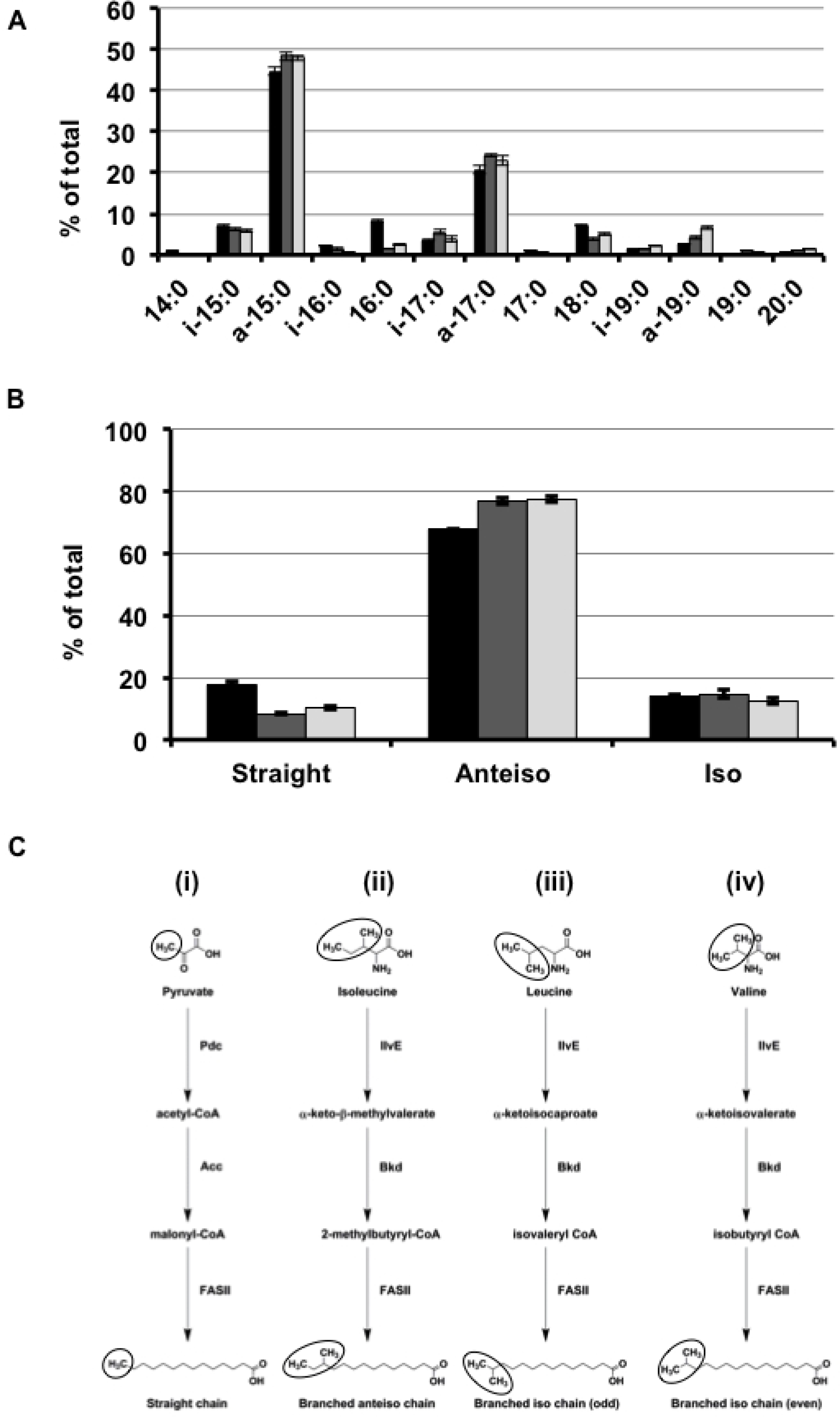
Fatty acid analysis from three different *S. aureus* strains. A) Total fatty acid composition of lipids from three *S. aureus* strains: BB270 (black), Newman (dark grey) and Mu50 (light grey). B) Relative percentage composition of straight, anteiso-branched and iso-branched chain fatty acids of lipids from three *S. aureus* strains: BB270 (black), Newman (dark grey) and Mu50 (light grey). C) Biosynthesis, including the FASII cycle, of the straight- and branched-chain fatty acids in *S. aureus*, illustrating the functional differences between the pathways. See S1 Table for a description of the genes involved in the pathways and their functions. Circled molecules show the origin for the structural differences between the different fatty acid types.

A simplified biosynthetic pathway of the straight- and branched-chain fatty acids identified by GC-MS in *S. aureus* is shown in Fig 1C. The specific genes that utilise the various molecular precursors to produce the straight-, branched-anteiso, branched-iso (odd chain) and branched-iso (even chain) fatty acids of these three *S. aureus* strains are listed in S1 Table, alongside other representative Gram-positive bacteria. As with most bacteria, straight-chain fatty acids are formed via pyruvate through the synthesis of malonyl-CoA that enters the normal type II fatty acid synthesis (FASII) pathway. The branched-chain fatty acids have amino acid precursors (valine, leucine and isoleucine) that, unlike pyruvate, are processed by an alternative suite of enzymes and enter the FASII pathway as specific branched-chain acyl-CoA substrates. Fig 1C highlights that it is the amino acid side chains that give the branched-chain fatty acids their ultimate structural identity.

### Lipid composition

#### Phosphatidylglycerols

Survey scans (nESI-MS) in negative ion mode of lipids extracted from each *S. aureus* strain revealed the profile of the most abundant lipid to be PG, with [M-H]^−^ ions ranging between m/z 679 – 777 (Fig 2). Submission of the parent masses to the LIPID MAPS database determined the PG lipids had a total fatty acid chain content of 29–36 carbons. This lipid profile was highly similar between each strain with only minor differences observed; the relative abundance of the greater molecular weight lipids (m/z 735 and higher) was lowest in BB270 (Fig 2A), greater in Newman (Fig 2B) and greatest in Mu50 (Fig 2C), and there were corresponding reductions in the PG species containing shorter fatty acid chains.

**Fig 2.**
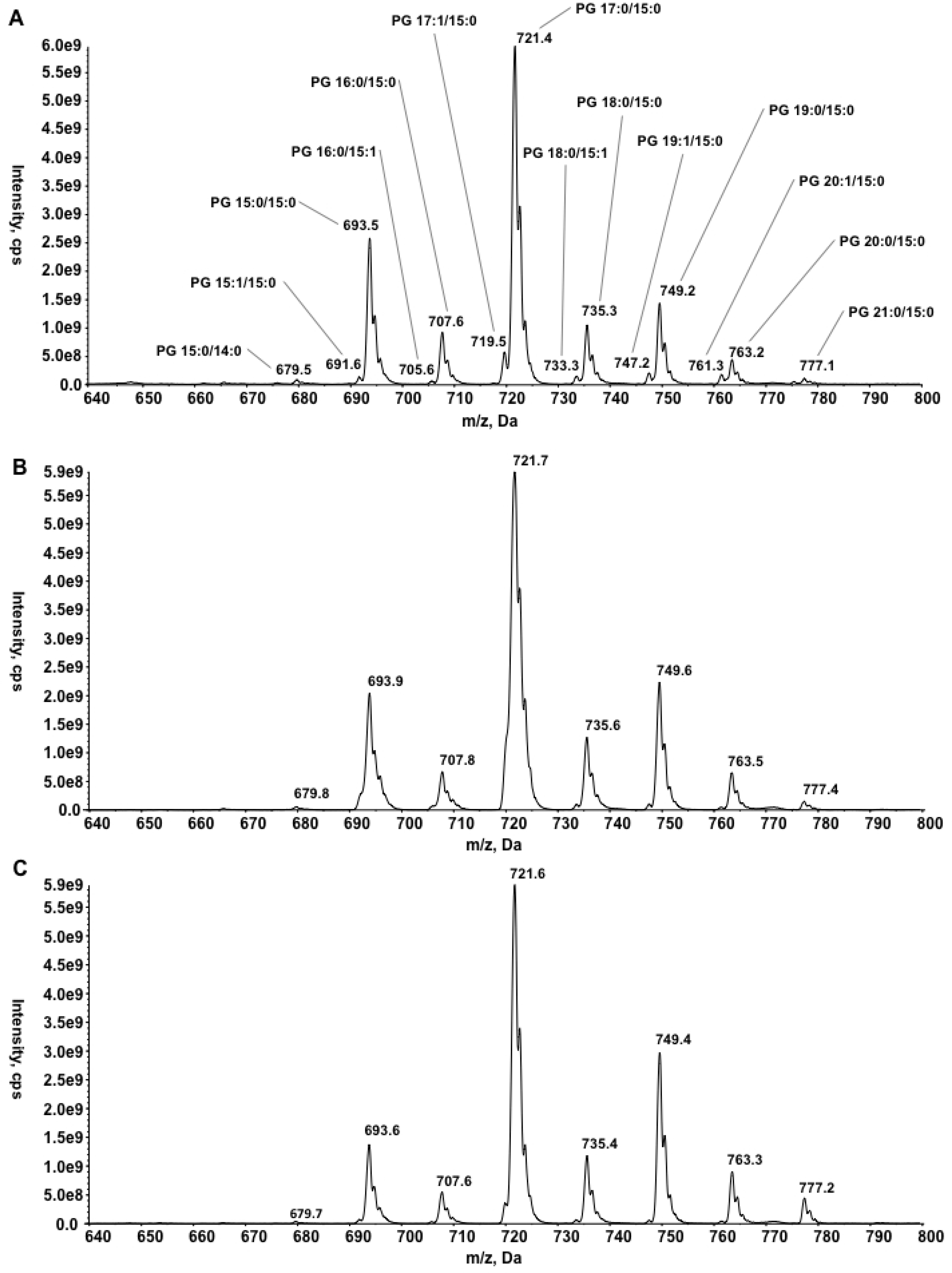
Survey scans in negative mode (640–800 m/z). Phosphatidylglycerol species of three *S. aureus* strains: A) BB270 B) Newman and C) Mu50.

In ESI-MS/MS, the established rule is that the more abundant daughter ions are those that result from loss of the acyl chain (as a neutral loss or as the ketene) from the 2 position (*sn*-2) of the glycerol backbone of the phospholipid [27]. Thus, parent ion fragmentation by nESI-MS/MS determined the specific composition and position of the acyl chains for each PG molecule (as an example, see the fragmentation spectrum of PG C32:0 in S1A Fig). The fatty acid composition of the principal isomers for each parent ion from the BB270 strain are noted in Fig. 2, while S2 Table also details the composition of the less abundant isomers. This revealed that it was always the shorter acyl chain of the two substituents that was found at the *sn*-2 position, most frequently a C15:0 fatty acid, with other saturated fatty acids in some of the minor PG species. This classification by nESI-MS/MS mirrored the total fatty acid abundance obtained by GC-MS, where C15:0 and C17:0 fatty acids predominated in the profiles (Fig 1A). As such, it is expected that the *sn*-1-linked C17:0 and *sn*-2-linked C15:0 fatty acids of the abundant PG molecule (m/z 721.4, C32:0) will predominantly be the anteiso-C17:0 and anteiso-C15:0 branched-chain versions identified to be the most abundant by GC-MS.

#### Diglucosyldiacylglycerols

ESI-MS survey scans in positive ion mode of lipids extracted from the three *S. aureus* strains revealed the presence of an abundant neutral lipid, diglucosyl-diacylglycerol (Glc_2_-DAG), with various [M+Na]^+^ ions ranging between 859–971 m/z (Fig 3). Glc_2_-DAG is the glycolipid that provides the membrane anchor for the principal cell wall polymer LTA [28]. Daughter ion fragmentation in positive mode confirmed each lipid was detected as a sodium adduct [M+Na]^+^ and total fatty acid chains ranged between 28–36 carbons (as an example, see the fragmentation spectrum in S2 Fig). Although all three strains had similar Glc_2_-DAG profiles with the dominant molecule again being C32:0, it is notable that, unlike BB270, Newman and Mu50 have Glc_2_-DAG lipids with fatty acid contents between 30–36 carbons with no shorter chain C28:0 or C29:0 detected. As Glc_2_-DAG lipids can be detected in negative mode as well as positive mode, positional assignments of the fatty acid chains were deduced from daughter ion spectra acquired from fragmentation of the lipids in both modes. It is important to note that in the fragmentation of molecular species in positive mode for glycolipids such as these, the more abundant daughter ions detected are those resulting from the loss of the acyl chain at the *sn*-1 rather than *sn*-2 position [29].

**Fig 3.**
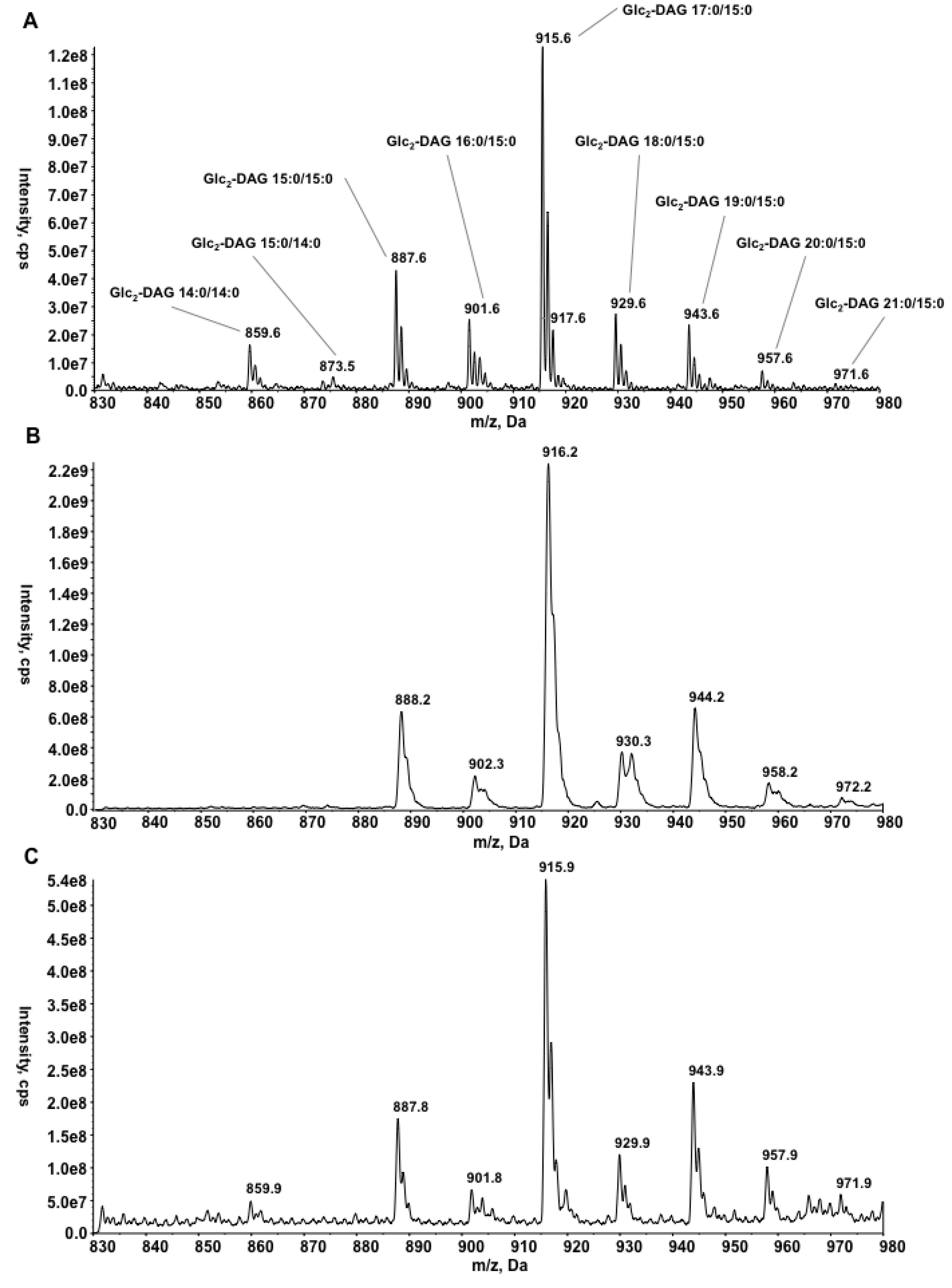
Survey scans in positive mode (830–980 m/z). Diglucosyl-diacylglycerol species of three *S. aureus* strains: A) BB270 B) Newman and C) Mu50.

#### Lysyl-phosphatidylglycerols

Neutral loss scans of 300 Da in positive ion mode of the three *S. aureus* lipid extracts revealed the profile of lysyl-PG, with peaks ranging 809–893 m/z (Fig 4). Daughter ion fragmentation in positive mode indicated each molecular species had a total fatty acid chain content ranging between 29-35 carbons (as an example, see fragmentation spectrum in S3 Fig) and showed that all three strains contained the same lipids, with only minor differences in their proportions. Like the PG that acts as the substrate for the formation of lysyl-PG, the C32:0 species is the dominant lipid in the profile, but a longer chain C36:0-containing lysyl-PG was not detected in any of the strains. Like Glc_2_-DAG, lysyl-PG can be detected in negative ion mode as well as positive and so the specific composition of each molecular species was established by daughter ion fragmentation in both modes. Again the acyl chain distribution was comparable to the composition of the other lipids (S2 Table) and mirrored the total fatty acid abundance obtained by GC-MS.

**Fig 4.**
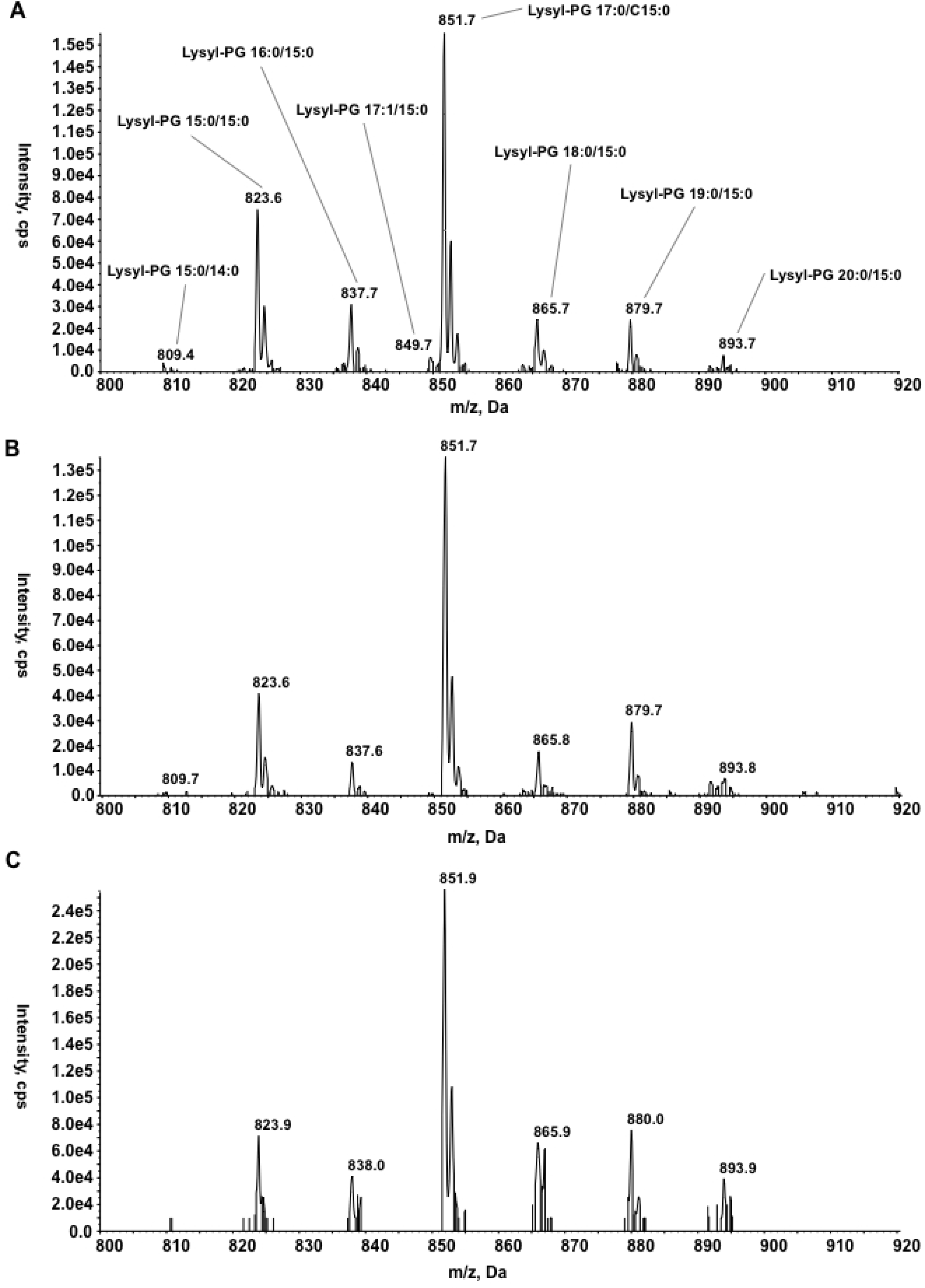
Neutral loss (300 Da) scans in positive mode (800-920 m/z). Lysyl-phosphatidylglycerol species of three *S. aureus* strains: A) BB270 B) Newman and C) Mu50.

#### Cardiolipins

The identification and characterisation of CL species in *S. aureus* can be difficult, mainly due to the minor abundance (~1%) present in exponential growth phase bacteria such as used in this present study [12]. Consequently, using the standard Bligh-Dyer extraction method resulted in limited detection of CL, and only in the BB270 and Newman strains (S4 Fig). The apparent lack of CL in the Mu50 strain might be due to the low efficiency of cardiolipin extraction from the cell membranes due to the thicker cell walls of the Mu50 strain [30]. In negative mode ESI-MS, the BB270 and Newman strains typically showed CL as [M-2H+Na]^−^ ions, though minor [M-H]^−^ species were also detected, a common observation when using this type of nESI-QTOF mass spectrometer [31]. Submission of the parent masses to the LIPIDMAPS database established that each molecular species had a total fatty acid chain content ranging 59-69 carbons and parent ion fragmentation by ESI-MS/MS determined the composition of the acyl chains (S2 Table). Predominantly, the fatty acid composition reflected the abundance of the PG species available from which to synthesise CL, though there is an absence of C19:0 and C20:0 due to the lack of detection of any higher molecular weight (i.e. greater than C69:0) species. Due to the difficulties in extraction and analysis, CL studies of *S. aureus* would benefit from targeted lipid analysis, a focus on stationary phase bacteria, and use of a modified Bligh-Dyer lipid extraction protocol utilising a pre-treatment step with lysostaphin [32], which is recommended for *S. aureus* strains with unusually thick cell walls [30].

#### Unsaturated lipids

Observable in some of the lipid profiles were a series of minor peaks that were 2 amu less than the equivalent major ions, suggesting the existence of lipids with unsaturation (specifically a single double bond) in some of the fatty acid chains. These ions are more prevalent in the PG survey scans and particularly prominent in the BB270 profile (Fig 2A). Fragmentation of these parent masses confirmed that a single fatty acid chain was unsaturated in each PG molecule (S2 table). Fragmentation also revealed that the double bond could be present in either an odd or even chain fatty acid in either the *sn*-1 or *sn*-2 position with no apparent specificity (S1B Fig). Though highly sensitive and analytical, nESI-MS/MS is unable to define the specific location of a double bond within the fatty acid chain. Single unsaturated fatty acids were detected also in some lysyl-PG and again were more prevalent in the BB270 strain, with fewer observed in the Newman strain and a complete absence in Mu50 (Fig 4). The fatty acid compositions of the unsaturated lysyl-PG species from BB270 were found to be very similar to the equivalent unsaturated PG species (S2 Table). Remarkably for all three strains, no unsaturation was detected in the Glc_2_-DAG lipids (Fig 3). As an alternative method to identify the presence of unsaturated fatty acids, precursor ion scanning was employed to detect parent PG species ions that contained fragments corresponding to unsaturated fatty acids (S5 Fig). For example, searching for a fragment ion of m/z 225 (C14:1) revealed no parent ions (this is in agreement with C14:1 not being detected in PG), while searching for parents of m/z 239 (C15:1) showed ions at m/z 691, 705, 719, 733, 747 and 761, in agreement with the daughter ion fragmentation of these parent ion species (S5A Fig). Conversely, m/z 267 (C17:1) was identified principally in a parent ion at m/z 719, the dominant molecular species indicated by fragmentation to contain the C17:1 fatty acid (S5B Fig). Furthermore, parent ion scanning revealed the existence of a small proportion of a doubly unsaturated (17:1/15:1) PG (m/z 717, S5 Fig and S2 Table).

The presence of unsaturated lipids in *S. aureus* is somewhat controversial, with a common assumption in the community that the biophysical role of unsaturated fatty acids is substituted by the abundant branched-chain fatty acids in Gram-positive bacteria [25]. Indeed, in two recent LC-MS studies on *S. aureus* lipids, one reported only saturated lipids to exist in Col and Newman strains [20], while the other on the N315 strain detected lipids containing unsaturated fatty acid chains (specifically C29:1 to C35:1 PG species) [21], though no comment was made on the existence of these molecular species. However, it is common to have minor variations in fatty acid contents and thus the profiles of bacterial lipidomes due to variations in growth phase, choice of strain and culture media, and indeed these earlier studies utilised brain-heart infusion media [20,21]. As the approved medium used for antimicrobial susceptibility testing, Müller-Hinton broth was selected for this present study to allow for future studies of the phenotyping of lipid and membrane changes induced by antimicrobial compounds.

To assess whether the detection of unsaturated fatty acids was specific to culture of *S. aureus* in Müller-Hinton broth, we analyzed the lipids of each of the three strains after culture in tryptic soy broth (TSB) and identified similar levels of unsaturated lipids from bacteria harvested under both culture conditions. S6 Fig shows the alternative profile of PG lipids from the Newman strain cultured in TSB (S6A Fig) in comparison to MH (S6B Fig). Fragmentation illustrated that the fatty acid composition of the PG species varied noticeably from that observed after culture in Müller-Hinton broth, with an increase in even-chain saturated FA relative to the branched-chain FA (S6 Fig). This observation concurs with a recent study that examined the fatty acid profiles of *S. aureus* after culture in diverse media including Müller-Hinton broth and TSB [26]. Unfortunately, GC-MS analysis of FAME from the three *S. aureus* strains failed to detect unsaturated fatty acids (Fig 1A). However, as the longer saturated-chain C21:0 fatty acid, observed in fragmentation of both PG and Glc_2_-DAG, was also absent from the FAME data, the lack of detection by the GC-MS is conceivably a question of sensitivity. While no obvious genes for the biosynthesis of unsaturated FA have been identified in the genomes of BB270, Newman and Mu50, a recent study showed that different strains of *S. aureus* cultured in bovine serum can readily acquire FA from the medium and incorporate these into their lipids such that monounsaturated FA can compose up to 30% of the total FA content [26]. Importantly, this shows that in a more physiologically relevant setting, *S. aureus* is flexible regarding the degree of fatty acid unsaturation present in lipids in its membranes.

Uptake and incorporation of exogenous fatty acids into *S. aureus* lipids (independent of *de novo* synthesis by FASII) is a recent discovery and occurs by a fatty acid kinase-dependent pathway [33]. Protonated fatty acids diffuse into the cell where they are bound by two fatty-acid binding proteins (FakB1/FakB2), with saturated fatty acids binding to FakB1 and unsaturated fatty acids to FAKB2 [34]. Using ATP FakA phosphorylates the fatty acid to acyl-phosphate (acyl-PO_4_), which can then be used as a substrate by the membrane-bound acyltransferase PlsY. *FakA* deletion prevents the incorporation of exogenous fatty acids into lipids and can lead to a stress response, increased resistance to antimicrobial peptides and decreased production of virulence factors [35]. Furthermore, when cultured in the presence of porcine liver extracts *S. aureus* bacteria with non-synonymous mutations in the malonyl CoA-acyl carrier protein transacylase (*fabD)* gene bypass the FASII pathway and incorporate long-chain polyunsaturated fatty acids such as C20:4 into their membranes [36]. Examination of a large panel of clinical isolates has indicated that it could be a common strategy of *S. aureus* to incorporate exogenous FA, overcoming its reliance on endogenous FA biosynthesis, and thus rendering ineffective FASII inhibitors such as the widely used biocide triclosan [37].

### Biosynthesis of phospholipids

All the *S. aureus* lipids characterised in this present study are synthesised by an interlinked biosynthetic pathway (Fig 5), and the respective genes that function in this pathway are presented for the three *S. aureus* strains analysed herein, and homologous genes from other Gram-positive bacteria, in S3 Table. Kuhn and colleagues describe in more detail the regulation of lipid biosynthesis in *S. aureus* [38].

**Figure 5:**
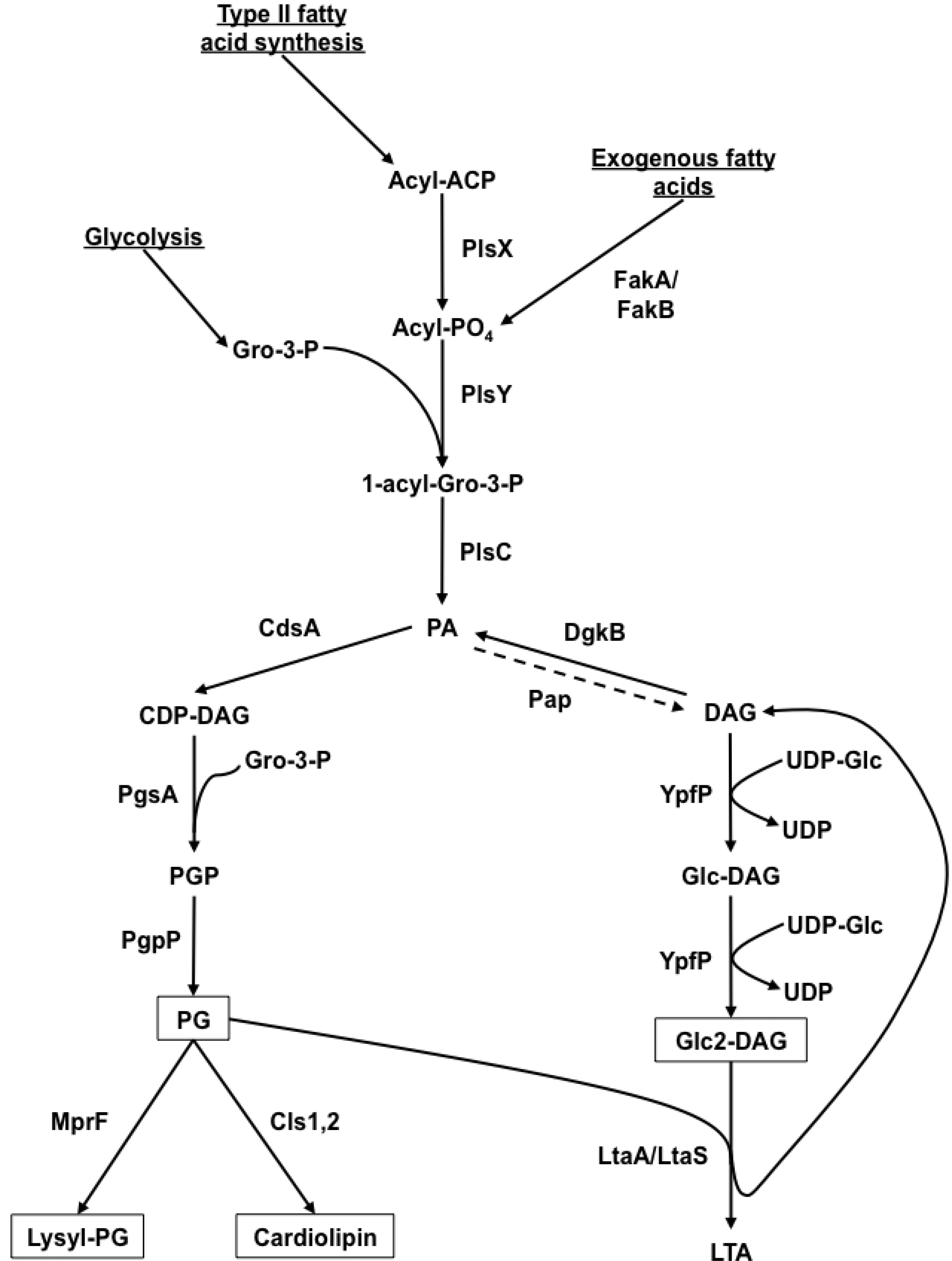
Lipid biosynthesis in *S. aureus*. Depicting the metabolites and proteins involved in the discrete pathways. The precursor molecules for lipid biosynthesis are underlined. Boxed molecules are those lipids that are readily detected in *S. aureus* cell membranes. A dotted line illustrates a theoretical recycling route from phosphatidic acid (PA) to diacylglycerol (DAG) yet to be confirmed experimentally. See S3 Table for a description of the genes involved in the pathways and their functions.

Briefly, synthesis of phospholipids in *S. aureus* is initiated by acylation of glycerol-3-phosphate (Gro-3-P) with fatty acids generated by the FASII pathway. Initially, the acyl-carrier protein substrates (acyl-ACP) produced by the FASII pathway are converted to acyl-PO_4_ [39] by the essential enzyme PlsX, whose deletion creates fatty acid auxotrophic S*. aureus* [33]. Subsequently, the similarly essential membrane-bound Gro-3-P acyltransferase PlsY uses the acyl-PO_4_ as substrate to form 1-acyl-Gro-3-P [40]. The second fatty acid is transferred to the *sn*-2 position of acyl-Gro-3-P from acyl-ACP by the membrane-bound PlsC, producing phosphatidic acid (PA).

Phospholipid synthesis continues with the production of cytidine diphosphate-diacylglycerol (CDP-DAG) by CdsA from PA using cytidine triphosphate. As a CDP-DAG-glycerol-3-phosphate 3-phosphatidyltransferase, PgsA then removes cytidine monophosphate and adds Gro-3-P to produce phosphatidylglycerolphosphate (PGP) [41]. Interestingly, mutations in *PgsA* have been identified in a daptomycin-non-susceptibility phenotype [42]. Significantly, the phosphatidylglycerophosphatase (PgpP) that removes the terminal phosphate to yield PG, the major phospholipid of *S. aureus*, remains to be identified [13]. Some of the PG pool is apportioned for further lipid synthesis, while some PG is synthesized to CL by the phospholipase D-like CL synthases Cls1 and Cls2, which fuse two PG molecules and release glycerol [43]. Cls1 seems to be necessary for CL synthesis only under conditions of acid stress, while Cls2 is the general bulk CL synthase utilized under normal culture conditions [32,44]. A relatively unusual but highly significant biological process is the aminoacylation of PG with L-lysine, forming lysyl-PG, by the multiple peptide resistance factor protein MprF that also translocates the cationic phospholipid from the inner to outer leaflet of the plasma membrane [45]. *MprF* is not essential, but deletion strains show much greater susceptibility to the detrimental effects of cationic antimicrobial peptides and antibiotics [46] and are attenuated in animal infection models [45].

Finally, as a constituent of the major cell wall component LTA [47], there is enormous demand for the Gro-P headgroup of PG, which requires the PG pool to be replenished up to three times per bacterial cell cycle [11, 48]. LTA is a polymeric glycerophosphate chain (n≈24) that is anchored in the plasma membrane by Glc_2_-DAG. As shown in Figure 6, this glycolipid anchor is synthesized with two consecutive glycosylations of DAG by the glucosyltransferase YpfP using UDP-glucose to form Glc_2_-DAG. Deletion of *YpfP* results in the absence of Glc_2_-DAG and as a consequence LTA anchors instead to the DAG precursor, resulting in drastically reduced viability when cultured under certain conditions, such as in 0.05 M phosphate buffer (pH 7.2) containing 0.85% NaCl at 37°C [28]. It is presumed that, as a putative glycolipid permease, LtaA translocates the Glc_2_-DAG to the outer leaflet of the plasma membrane, where the membrane-bound LtaS transfers 20–40 glycerophosphate molecules from PG lipids to the elongating glycolipid anchor [49]. The DAG generated from this transfer is recycled by the DAG kinase DgkB that phosphorylates it to PA, which can be re-utilized for further PG synthesis [50]. Unsurprisingly, deletion of *LtaS* produces a *S. aureus* cell lacking in LTA [49] and that is unable to grow unless compensatory mutations are present to increase the pool of the secondary messenger c-di-AMP [51].

## Conclusions

In conclusion, this present study reports the application of a rapid, non-targeted nESI-MS/MS lipidomic method allowing characterisation of three representative *S. aureus* strains. This technology gives a high sensitivity of detection and provides a more complete description of lipid composition in comparison to previous studies, including the positional assignments of fatty acids. This enhanced sensitivity has enabled the detection of comparatively minor differences in the lipid profiles of the *S. aureus* strains under investigation. As a result it can be routinely employed to compare *S. aureus* lipid profiles, which may assist with determining the role of membrane lipids in the development of resistance to membrane-active antimicrobial agents, and could be developed even further to diagnose drug-resistant strains in the clinic.

## Acknowledgments

We gratefully acknowledge Ms. Leigh-Ann Booth (University of St Andrews) for assisting in the assignment of lipid compositions from fragmentation spectra.

## Supporting Information

**S1 Figure. A) Daughter ion spectra in negative mode of C32:0 PG (m/z 721 [M-H]^−^) and B) Daughter ion spectra of C32:1 PG (m/z 719 [M-H]^−^).** The insets show representative structures of the lipids (with straight chain fatty acids for simplicity) detailing some of the characteristic daughter ion fragments.

**S2 Figure. Daughter ion spectra in positive mode of C32:0 Glc_2_-DAG (m/z 915 [M+Na]^+^).** The inset shows a representative structure of the lipid (with straight chain fatty acids for simplicity) detailing some of the characteristic daughter ion fragments. A common ESI-MS contaminant in positive mode is highlighted by *.

**S3 Figure. Daughter ion spectra in negative mode of major C32:0 lysyl-PG species (m/z 849 [M-H]^−^).** The inset shows a representative structure of the lipid (with straight chain fatty acids for simplicity) detailing some of the characteristic daughter ion fragments.

**S4 Figure. Survey scans in negative mode (1300-1450 m/z) of cardiolipin in *S. aureus* strains A) BB270 and B) Newman.** Both major [M-2H+Na]^−^ and minor [M-H]^−^ ions are highlighted alongside [2(M-H)+Na]^−^ dimers of abundant PG lipids (*).

**S5 Figure. Precursor ion scans in negative mode of *S. aureus* BB270 showing phosphatidylglycerol species as A) Parent ions of m/z 239 and B) Parent ions of m/z 267.** The fatty acid compositions for the alternative mass isomers identified in the other parent ion scan are written in grey.

**S6 Figure. Survey scan in negative mode (640-800 m/z) of phosphatidylglycerol species from A) *S. aureus* Newman cultured in Tryptic Soy Broth and for comparison B) *S. aureus* Newman cultured in Müller-Hinton Broth.** Note that the survey scan shown here in panel B is identical to that displayed in Figure 2 panel B.

**S1 Table. Genes with putative functions in fatty acid biosynthesis**. Specific genes (as illustrated in figure 1C) are listed from the genomes of *S. aureus* strains NCTC 8325 (parent strain of BB270), Newman and Mu50. Included (though not illustrated in Figure 1C) are the type II fatty acid biosynthesis genes (FASII) designated ‘Fab’. Homologous genes from other Gram-positive bacteria, *Streptococcus pneumoniae* R6 and *Bacillus subtilis* 168, are shown for comparison. Genes in bold have been designated essential based on genetic disruption in *S. aureus* NCTC8325 [52], *B. subtilis* 168 [53,54] and *S. pneumoniae* R6 [55].

**S2 Table: All characterised lipid species detailing composition and acyl chain position**. Based upon fragmentations of lipids from all three *S. aureus* strains: BB270, Newman and Mu50.

**S3 Table. Genes with putative functions in lipid biosynthesis.** Specific genes (as illustrated in Figure 5) are listed from the genomes of *S. aureus* strains NCTC 8325 (parent strain of BB270), Newman and Mu50. Homologous genes from other Gram-positive bacteria, *Streptococcus pneumoniae* (strain R6) and *Bacillus subtilis* (strain 168), are shown for comparison. Genes in bold have been designated essential based on transposon mutagenesis analysis in *S. aureus* NCTC8325 [52], *B. subtilis* 168 [53,54] and *S. pneumoniae* R6 [55,56]. *Note that *S. pneumoniae* LTA has a distinct chemical composition to that of *S. aureus* LTA [57] and thus the *LtaA* and *LtaS* genes are unrelated.

